# Variational graph encoders: a surprisingly effective generalist algorithm for holistic computer-aided drug design

**DOI:** 10.1101/2023.01.11.523575

**Authors:** Hilbert Lam Yuen In, Robbe Pincket, Hao Han, Xing Er Ong, Zechen Wang, Weifeng Li, Jamie Hinks, Liangzhen Zheng, Yanjie Wei, Yuguang Mu

**Affiliations:** School of Biological Sciences, Nanyang Technological University, Singapore, 60 Nanyang Dr, Singapore 637551; A*STAR Skin Research Labs, Agency of Science, Technology and Research, Singapore, 11 Mandalay Rd, #17-01, Singapore 308232; School of Physics, Shandong University, Jinan, Shandong 250100, China; Singapore Centre for Environmental Life Sciences Engineering, Nanyang Technological University, Singapore, 60 Nanyang Dr, Singapore 637551; Shenzhen Institute of Advanced Technology, Chinese Academy of Sciences, Shenzhen, Guangdong 518055, China; Shanghai Zelixir Biotech, Shanghai 200030, China

## Abstract

While there has been significant progress in molecular property prediction in computer-aided drug design, there is a critical need to have fast and accurate models. Many of the currently available methods are mostly specialists in predicting specific properties, leading to the use of many models side-by-side that lead to impossibly high computational overheads for the common researcher. Henceforth, the authors propose a single, generalist unified model exploiting graph convolutional variational encoders that can simultaneously predict multiple properties such as absorption, distribution, metabolism, excretion and toxicity (ADMET), target-specific docking score prediction and drug-drug interactions. Considerably, the use of this method allows for state-of-the-art virtual screening with an acceleration advantage of up to two orders of magnitude. The minimisation of a graph variational encoder’s latent space also allows for accelerated development of specific drugs for targets with Pareto optimality principles considered, and has the added advantage of explainability.

## 2. Introduction

As there is an estimated chemical space of 10^63^ molecules, comprising of the combination of 30 carbon, nitrogen, oxygen, and sulfur atoms in different arrangements^1^, the possibility of drug discovery is endless. However, high attrition rates in drug discovery^2^ is an overarching problem in biomedical sciences, with many “casualties” incurred in the multistep process of gaining regulatory approval for a drug. It is estimated, as of 2020, that the developmental cost of each drug approved by the United States Food and Drug Administration (FDA) costs an average of USD 1.3 billion^3^. As a result, computer-aided drug design (CADD) is an important field in which an initial screen of molecules is conducted and further optimisation can be performed. Good leads discovered at the initial stages are hence crucial to the drug discovery process^4^.

However, major problems exist in CADD, in which the authors flag three: one - the high computational costs involved. Although many CADD tools are largely democratised and freely accessible, computational power in the leagues of high-performance computing (HPC) supercomputers required for CADD^5^ is unfortunately expensive and unattainable to many researchers. Two - the likelihood of a drug making it to the stage of human consumption is not solely based on efficacy as a treatment against its intended disease. 90% of drugs fail to make it past clinical trials^6^. Many drug properties such as absorption, distribution, metabolism, excretion and toxicity (ADMET), drug-drug interactions (DDI)^7^ and side effects largely influence the success of a drug^8^. Three - current techniques in CADD typically involve the use of many specialised models, with each model predicting a specific chemical property^9^. When stacked with many models, computational cost required rises exponentially. Furthermore, there is a current trend of small molecule drug discovery in which these factors have a stronger weight than before, in which only drug efficacy and binding was considered^10,11^. With successful drugs that make it into the market typically having carefully balanced traits, this is also one factor for consideration.

Therefore, to address the stated problems in CADD, the authors propose a “variational graph encoder” (Fig. 1) - a convolutional graph neural network model encompassing elements from the variational autoencoder^12^ that is trained to predict simple descriptors of molecules and binary molecular fingerprints (FP) instead of reconstructing the input. From the intermediate mathematical representation of the variational graph encoder - also known as the latent space, surrogate models can then be trained to predict more complicated properties. Previous work involved utilisation of latent space include sampling in variational autoencoders to generate potent and selective RIPK1 inhibitors^13^ and BRAF inhibitor development^14^.

**Figure 1.**
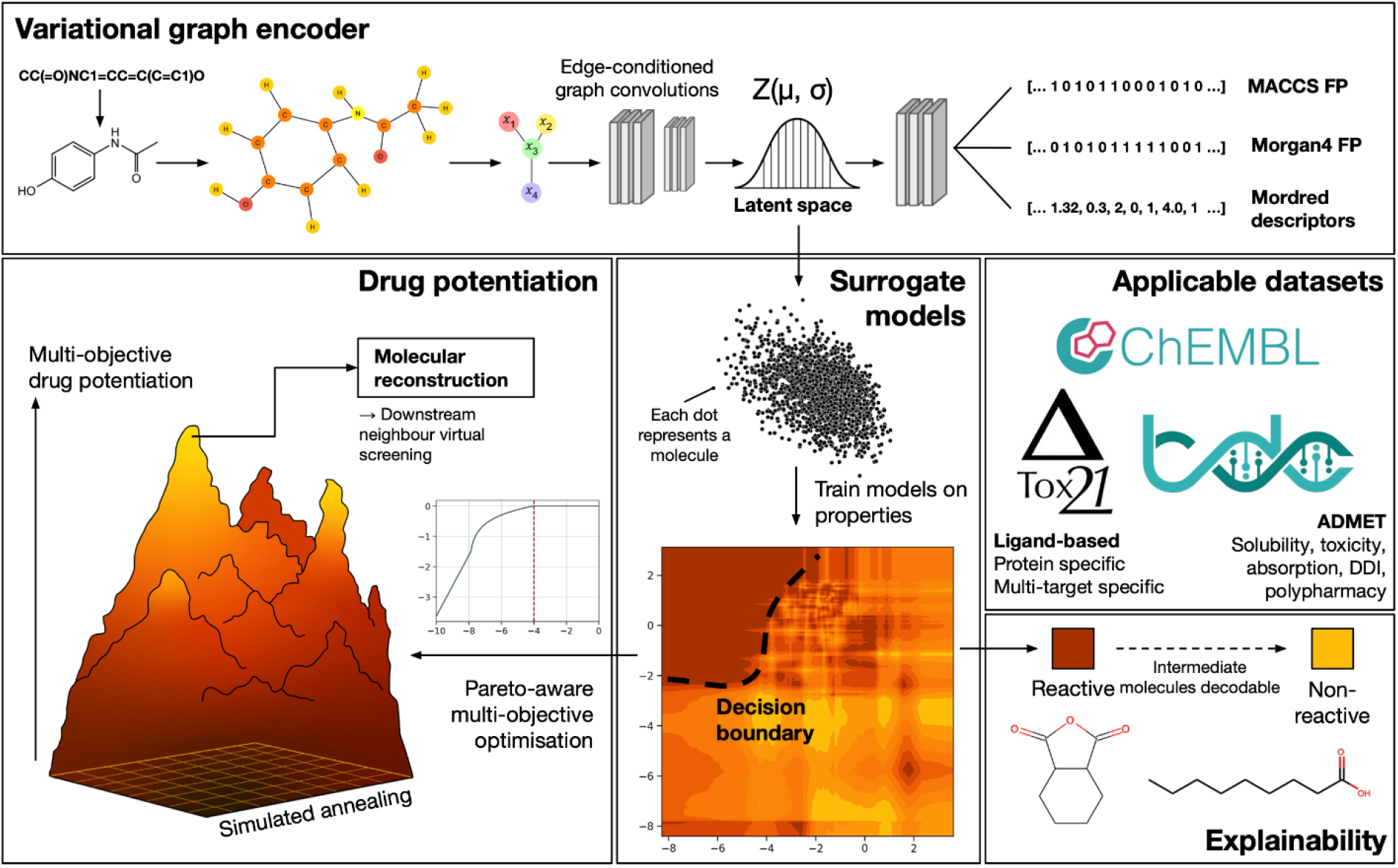
Molecules are encoded into a graph format, which is then passed through an autoencoder, with intermediate mathematical latent space is used for property prediction through surrogate models. SMILES strings are converted into molecular graphs, which are then convolved through edge-conditioned graph convolutional layers. Global sum pooling is then applied, and a latent space of 64 is achieved. When the variational graph encoder is trained, the model goes on to produce MACCS, Morgan4 FPs and Mordred descriptors. The latent space is separately extracted to train surrogate models, and decision boundaries can be plotted. Molecules can then be potentiated with respect to Pareto optimality principles with multiple objectives and reconstructed. Molecules at different intersections of decision boundaries can also be reconstructed to provide explainability as to why one molecule fits into one category but not another. Applicable datasets in which surrogate models can be trained on include datasets from ChEMBL, Tox21 and TDCommons databases, with ADMET properties, drug interaction/side effects being predicted. Ligand-based drug discovery is also possible with the use of surrogate models that are protein-specific, or are organism specific, such as prediction of results from multi-factorial coupled assays.

Methodology-wise, the previous work encoding and decoding SMILES strings in recurrent neural networks unlocked the possibilities of latent space optimisation and prediction for molecules^15^. Subsequent work using variational autoencoder neural networks with graph features and adjacency matrices concatenated also showed promising results^16^. However, the latter limits the size of the molecule that can be fitted into or generated by the autoencoder, whereas the former is susceptible to having multiple SMILES strings encoding for the same molecule. Both presented methods do not involve any node-level convolution. Our solution to this is to use the edge-conditioned graph convolutional neural networks^17^, which actively deciphers the connections and neighbouring atoms in each molecule through convolutions. The encoding of fingerprints (FPs) and chemical descriptors instead of a traditional autoencoder also allows for molecules of any size to be encoded whilst maintaining the bond and connectivity information. As a result, with these limitations overcome, accurate and explainable models predicting datasets from the Tox21^18^, Therapeutic Data Commons (TDCommons)^19^ and ChEMBL^20^ databases could be achieved in tandem with target-specific score functions in virtual screening^21^.

With a greatly increased overall diversity and accuracy of surrogate models, more challenging multi-objective optimisation could then be conducted using Pareto optimisation principles. When applied in tandem with structure-based virtual screening augmented with the proposed model, an acceleration of up to 2 orders of magnitude can be observed in the initial screening. This allows for the screening of massive numbers of molecules that was previously impossible. The authors hence anticipate that the use of such methodology will shift the pendulum from CADD to computer-aided drug engineering (CADE).

## 3. Results

### 3.1 Graph variational encoder shows high capability in deriving molecular descriptors while maintaining an evenly-distributed latent space

Overall, when the graph variational autoencoder was trained with molecular graphs derived from SMILES strings, the model generally proved an median accuracy of > 90% on Morgan4 and MACCS FPs, with the AUROC and AUPRC metrics being lower for Morgan4 FP than MACCS. The latents were also noted as being approximately normally distributed across 0 on all dimensions. The addition and subsequent increment of the KL loss at epoch 11 caused a significant dip in overall performance, however this was recuperated with further training (Fig. 1a). Without the addition of KL loss, the latent space showed an overall larger deviation from the Gaussian distribution and with larger individual latent space values. Mordred fingerprints performed reasonably well, with the majority of MAEs < 1 for each descriptor (Fig. 1d). The usage of a cis-trans-aware model was also performed with extra graph edges and did not show significant benefit in the FP classifier or Mordred descriptor regression results, hence the simpler model without added functionality was chosen. Clustered ZINC molecules using the Morgan2 fingerprint with 256 bits overall showed visual molecular similarity (Supplementary Fig. 2), with molecules displaying very similar backbones. Reconstruction of non-centroid molecules generally showed higher fidelity with identification through Euclidean distance of latent space than through FP matching, with the number of rings and substituents better preserved, although in both cases the correct substituent types were reconstructed (Supplementary Fig. 3).

### 3.2 Surrogate models accomplish similar accuracies when compared with specialist models on existing datasets in ADMET predictions, show strong generalisability for multi-class, multi-property problems, and can be applied to niche datasets

Best classification was achieved with extra trees classifiers compared to other models tested. The model performed well for the blood brain barrier dataset and human ether-a-go-go (hERG) datasets (Fig. 3a). Median AUROC and AUPRC and their respective median errors for the TDCommons classification dataset is 0.870 ± 0.021 and 0.891 ± 0.020 respectively. Performance was also topped for androgen and oestrogen receptor antagonists and agonists in the Tox21 dataset (Fig. 3b) and generally were 5th and above for all other datasets when AUROC was compared. Tox21-trained surrogate models had a median AUROC and AUPRC and their respective median errors of 0.854 ± 0.021 and 0.985 ± 0.003 respectively (Supplementary File 1). The DDI dataset, which is a multi-class problem, also showed AUPRC scores of > 0.975 (Fig. 3c). The TWOSIDES polypharmacy set showed overall low AUPRC scores when raw labels were used, however when reclassified into 26 categories of side effects using the ICD-11 as a guide, the score increased to an overall AUPRC of > 0.75 (Fig. 3d). Regression datasets were similarly applicable, with good prediction in the LD_50_ dataset and intestinal epithelial cell permeability (Caco2 dataset) from the TDCommons database, with Spearman’s rhos exceeding 0.7 in both instances. Other datasets tested also showed similarly strong correlations (Supplementary File 1). Specialised, niche datasets as extracted from ChEMBL likewise showed > 0.5 in Spearman’s rho when predicting the WHO cytotoxicity dataset [ChEMBL2093836] and fraction of unbound blood protein after IV drug administration [ChEMBL1614672] (Fig. 3f). The latent space also shows human-understandable decision boundaries for both classification (Fig. 3g) and regression tasks (Fig. 3h). Furthermore, in plotting the decision boundaries for the P-glycoprotein set, only two dimensions could be used, and the AUROC showed a dip of around 0.16. This provides further evidence that the use of more dimensions of the latent space provides more molecular description that greatly improves prediction quality. The TDCommons leaderboard uses a “stricter” scaffold split instead of a random split as shown in this work’s figures. Despite this, the surrogate models trained on the variational graph encoder’s latent space also topped the leaderboard in the datasets CYP2C9 and CYP2D6, with AMES and blood brain barrier penetration coming in second. Similarly, for regression problems, half life prediction and volume of distribution at steady state came in second using a support vector regression model (Supplementary File 1).

**Figure 2.**
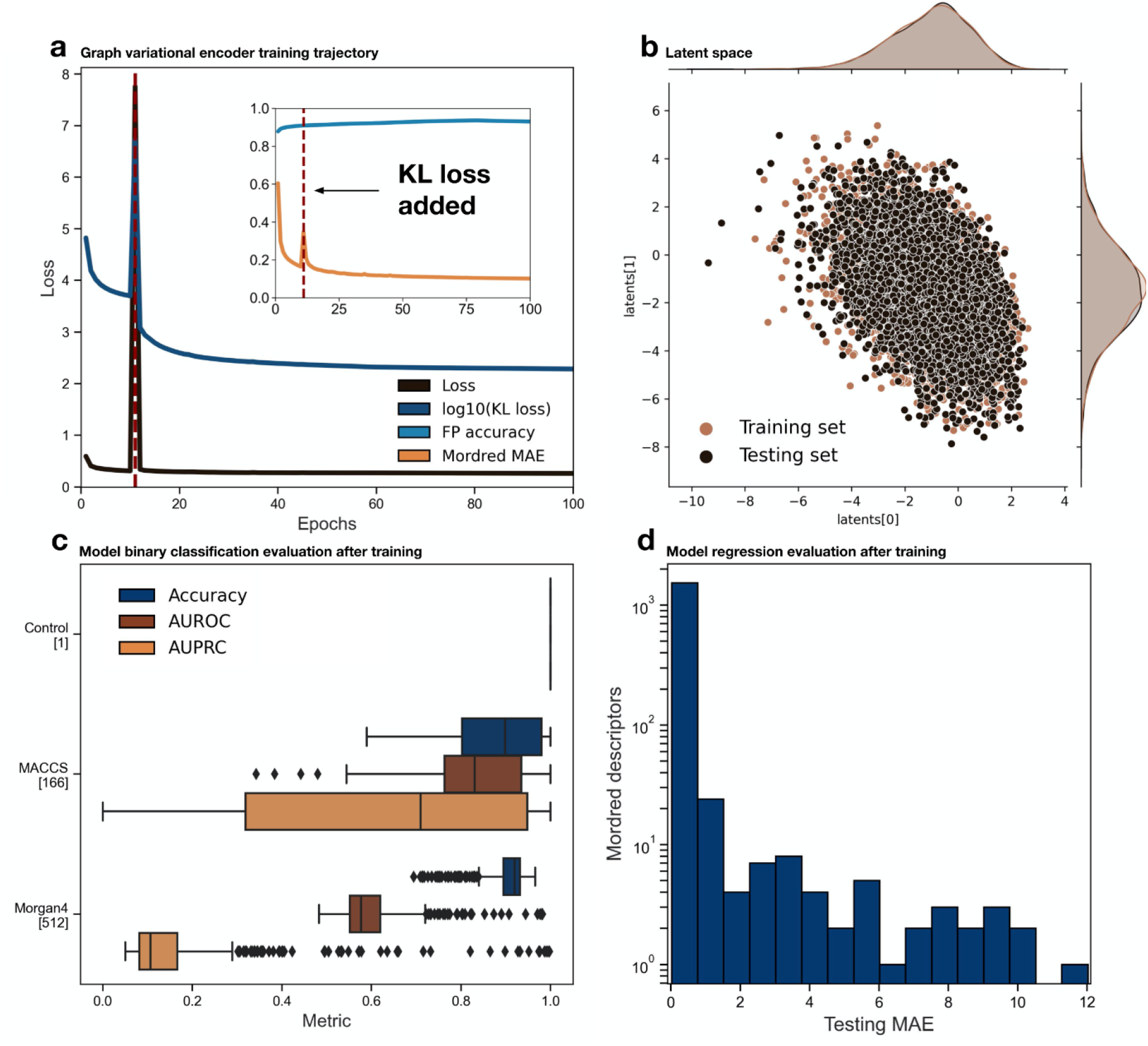
Variational graph encoder showed high accuracy in deriving fingerprints and other molecular descriptors whilst maintaining a Gaussian-distributed latent space. (a) Training loss, KL loss, and associated metrics for the model which was trained for 100 epochs on 650,000 unique molecules clustered from the ZINC database. (b) Visual of first and second dimension of the latent space shows an overall similarity in distribution in the training and testing set which were randomly split, showing 5,000 samples from each group. (c) Accuracy, AUROC and AUPRC are expectedly the highest in the control set, followed by the MACCS FP and Morgan4 circular FP. The results are derived from the 50,000-molecule testing set in which the model was not trained on. (d) Histogram of MAE of Mordred descriptors shows an overall ability of the model to integrate FP predictions, molecular graphs and associated chemical properties.

**Figure 3.**
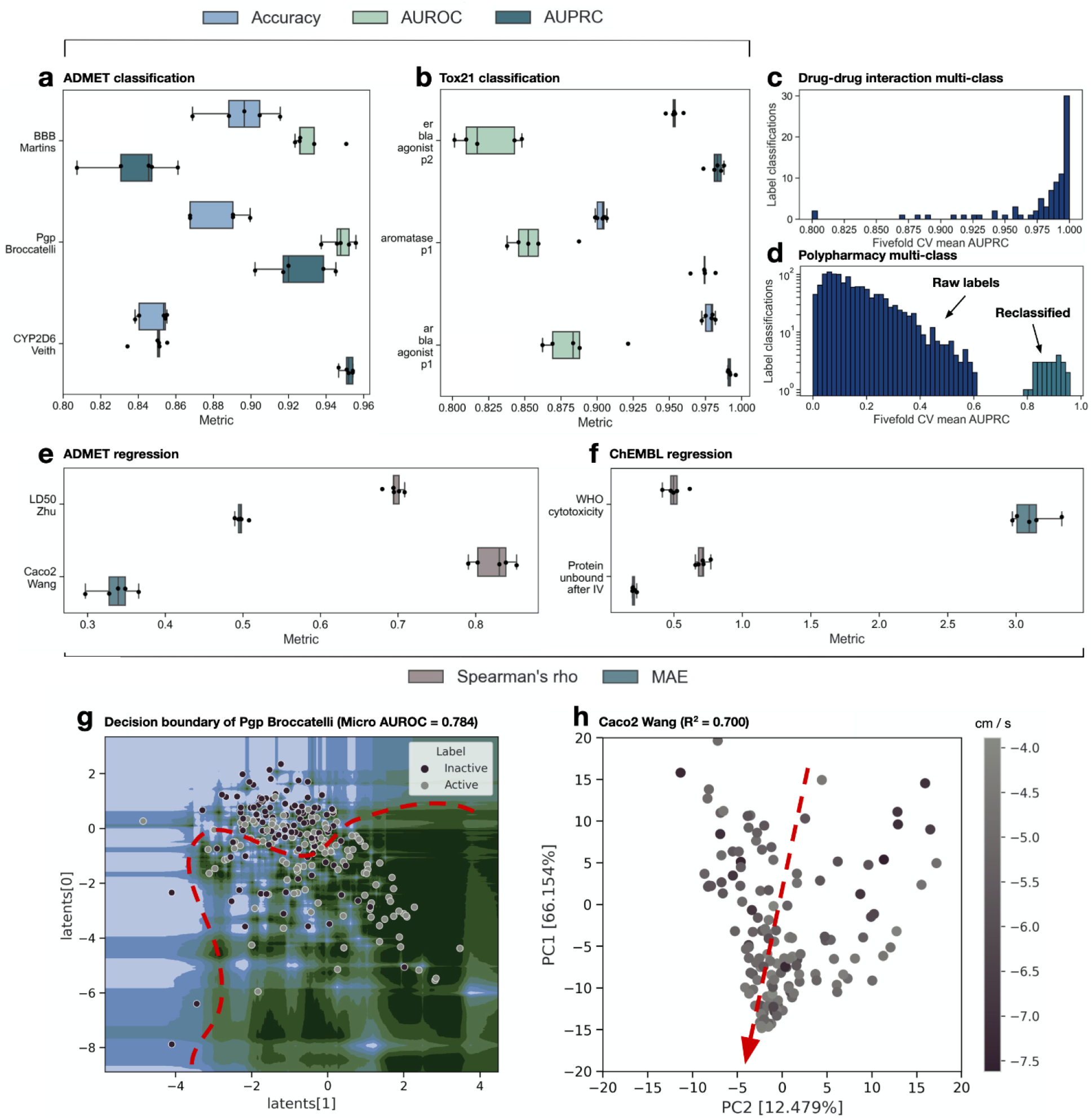
Surrogate models exploiting the variational graph encoder’s latent space can accurately predict single- and multi-classification problems, and regression problems for common datasets, even if the data is skewed. (a) Fivefold cross-validated ADMET classifications of blood brain barrier penetration, P-glycoprotein inhibition of CYP2D6 liver enzyme. (b) Fivefold validations Tox-21 datasets of oestrogen receptor alpha, aromatase, and androgen receptor signalling pathway. (c) Histogram plot of fivefold AUPRC results from multi-class problem of drug-drug interactions with a total of 84 classes. (d) Histogram plot of fivefold AUPRC results from multi-class problem of polypharmacy side effects from the TWOSIDES dataset. Blue indicates the raw labels, whereas cyan indicates the ICD-11 reclassified labels. (e) Fivefold ADMET regression consisting of MAE and Spearman’s rho consisting of the LD50 dataset and Caco2 dataset for molecule epithelial drug penetration. (f) Fivefold MAE and Spearman’s rho of ChEMBL regression datasets for cytotoxicity and ratio of protein unbound to plasma after intravenous administration. (g) Decision boundary of an extra trees classifier model trained on the P-glycoprotein dataset with only two dimensions of latent space from the variational graph encoder, with green indicating regions where the surrogate model predicts active binding and blue indicating inactive binding, with stronger colours indicating higher confidence. (h) Regression and associated gradients plotted with PCoA on the latent space for the Caco2 dataset. All box and whisker plots shown reflect the standard 1.5 IQR fence with the box encompassing 50% of the median data.

### 3.3 Ligand-based drug discovery is possible using variational graph encoders, with surrogate models taking near-negligible amounts of time to train

Surrogate models training on the 64-dimension latent space generally did not exceed one minute to train for 10,000 data points using SVMs. The decoding of a SMILES string into the corresponding latent space also took less than 100 ms per molecule (Fig. 4f). Adding the time taken to obtain the latent space and the near-negligible time to predict ligand properties in surrogate models, this method provides an acceleration of 1.5 to 2 orders of magnitude when compared to existing score functions. The models tested were also performant, being able to tackle both classification (Fig. 4a - b) and regression problems (Fig. 4c - d) on existing score functions. Classification models showed high AUROC scores for influenza A spike protein from the ChEMBL database (ID: ChEMBL1614236) and SARS-CoV-2 protease from the TDCommons database (> 0.8 for both), whereas regression models for Vina scores, Gnina, OnionNet-SFCT, RTMScore and DeepRMSD gave an average Spearman’s rho values of > 0.5 in all tested instances (Fig. 4e), with Vina scores showing the highest correlation (Fig. 4c, e). In multimodal distributions, outlier values for Vina score functions were not as well modelled, with multimodal distributions showing overall poor R^2^ values (Supplementary Fig. 4), whilst maintaining a Spearman’s rho of > 0.6. This indicates that the surrogate model excels at predicting rankings in a particular score function, but not in predicting the exact scores directly.

**Figure 4.**
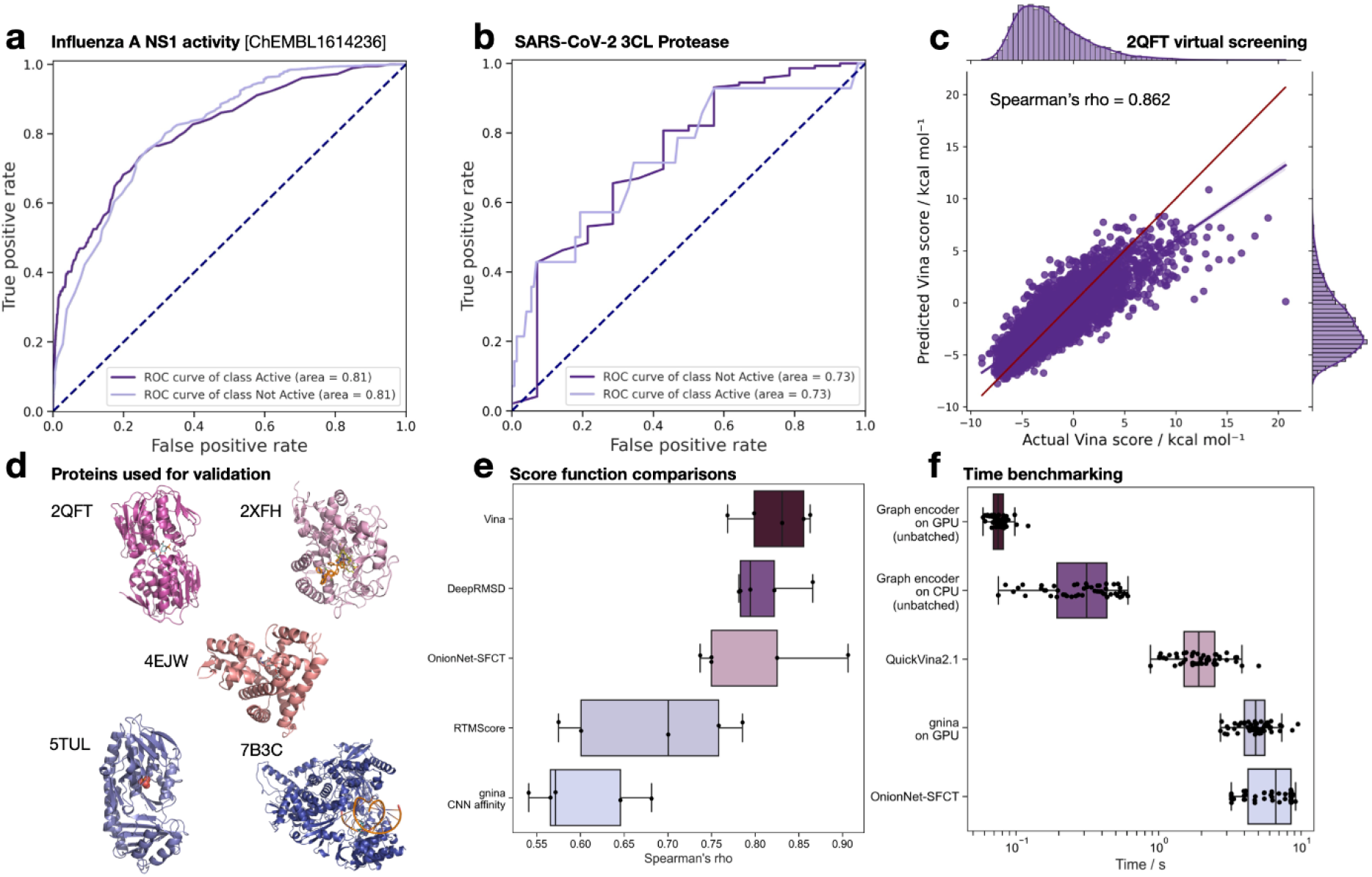
Ligand-based drug discovery is doable with latent space-trained surrogate models, with significant speedup. ChemEMBL and TDCommons datasets consisting (a) influenza spike protein activity and (b) SARS-CoV-2 protease were placed into an extra trees classifier and evaluated on a 20% split test set. (c) NuSVR surrogate model was trained to predict Vina score from the latent space of 45,000 molecules and evaluated on a test set of 5,000, with red line indicating perfect fit and purple line indicating the line of best fit. (d) Proteins with PDB IDs 2QFT, 2XFH, 4EJW, 5TUL and 7B3C and their native ligands which were used for the docking and scoring. (e) Fivefold regression of score functions on five chosen proteins using the NuSVR algorithm, with a hard cutoff of 100,000 iterations for Gnina due to the long convergence time. (f) Speed at which a molecule’s latent space is derived versus commonly used score functions.

### 3.4 Optimisation of latent space allows for molecule reconstruction through latent space comparison

Simulated annealing optimisation showed that multi-objective optimisation is possible using Pareto techniques (Fig. 5b - d). Ten unique parameters optimised all reached desired or near-desired set points for the *E. coli* EPSP synthase 2QFT protein: Vina score, aqueous solubility, five separate CYP P450 enzyme non-inhibition, human ether-a-go-go (hERG) non-inhibition, lack of mutagenic properties (AMES), and passability in the ClinTox dataset (Fig. 5c). The molecular trajectory clearly shows the addition of more oxygen-based substituents, increasing overall solubility. Simulated annealing also showed thorough exploration of the latent space in the initial iterations (Fig 5a). Redocking of the final molecule obtained from latent space optimisation yielded a binding score of −6.7 kcal/mol in Vina score (Fig. 5i), which is an improvement over the initial ligand’s docking score of −2.0 kcal/mol. The reconstructed intermediate molecules from the trajectory also show an increasing trend of Vina score, moving from −2.0 kcal/mol to −3.0 kcal/mol to −4.8 kcal/mol and lastly to −6.7 kcal/mol. Overall, optima was achieved in approximately 5,000 iterations of simulated annealing, with little further improvement in the reward scores for later iterations (Fig. 5g). All objectives were fulfilled except AMES mutagenicity, which scored 1.4% higher than the set target of 40% confidence of mutagen and 60% confidence of non-mutagen (Fig. 5e).

**Figure 5.**
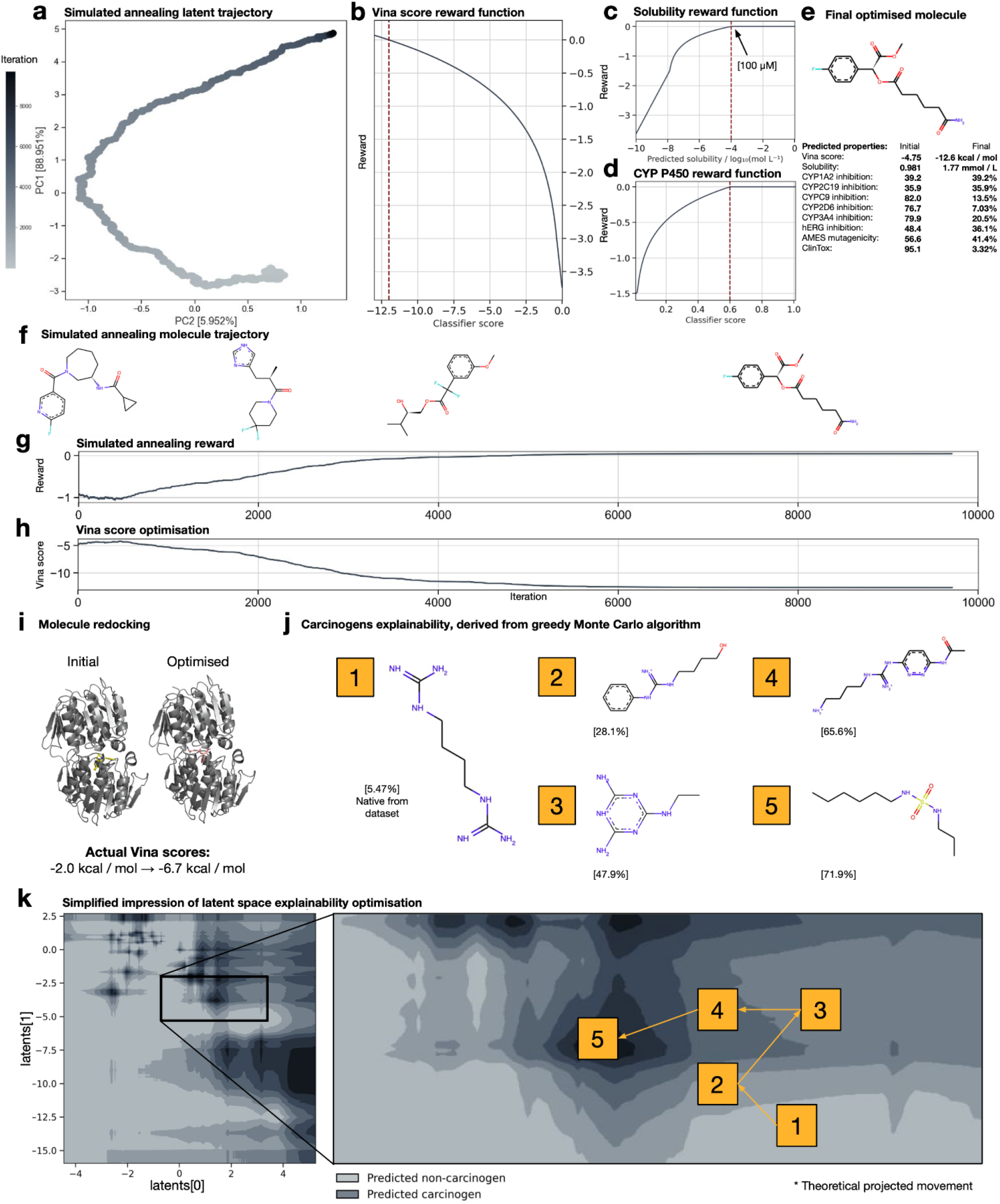
Desired molecular properties can be engineered with surrogate model optimisation, with explainability as to how one molecule is preferred over another in property prediction. Random latent space potentiation was performed with simulated annealing on the 2QFT *E. coli* holo-form EPSP synthase protein catalytic site, with the surrogate Vina score function model being a NuSVR trained on 45,000 randomly docked molecules from the ChemBridge core and express libraries. (a) The trajectory of the simulated annealing process. (b, c, d) The Pareto-optimality reward function for Vina score, solubility, and CYP P450 non-inhibitor. (e) The final output molecule from potentiation, with initial and final predicted properties that were optimised in accordance to the reward functions. (f) Decoded molecular trajectory over the steps. (g) Simulated annealing reward across iterations. (h) Vina score throughout iterations, with lower score indicating better binding. (i) Docked pose of reconstructed initial and final molecules with respective ground Vina scores. (j) Trajectory of non-carcinogenic molecule 1,4-diguanidinobutane and carcinogenic property confidence score when the latent space was potentiated using a greedy Monte Carlo algorithm, with formation of previously deemed potential carcinogenic sulfamide group in part 5. (k) Impression of latent space optimisation using the greedy Monte Carlo algorithm. Molecular reconstruction was performed using latent space comparison to the clustered ZINC dataset, with the molecule having the lowest Euclidean distance in latent space returned.

### 3.5 Latent space provides good explainability through the use of decision boundaries

Drug explainability was indicated in a non-carcinogenic molecule in the carcinogens dataset by Lagunin *et al*., 2009^22^, when the greedy Monte Carlo algorithm deployed slowly picked increasingly predicted carcinogenic properties, as illustrated in the decision boundary impression (Fig 5k). Furthermore, the trajectory shown (Fig. 5j) shows the formation of a sulfamide functional group, in which a superset group sulfonamide has previously shown to significantly increase tumour formation in mice^23^ and can cause follicular cell adenocarcinomas in rats^24^.

## 4. Discussion

The variational graph encoder and downstream surrogate models showed near-to-current state-of-the-art performance with significantly increased speed and could be used to expedite the drug discovery process, with latent space drug design being able to be performed on a personal laptop. Moreover, the ability of the model to encode a theoretically infinite chemical space into a continuous mathematical one proves benefit in a paradigm shift from CADD to CADE, allowing for fully *de novo* drug development, given limited empirical information about a protein target. Previous models that have shown highest classification ability for polypharmacy interactions required knowledge of multiple protein targets for drug-protein interactions^25^, with the proposed model able to do it *ab initio*.

Furthermore, the usage of graph convolutional neural networks allows for molecules to be embedded in an intuitive format. Previous work published on deep learning models have been done either on SMILES using a recurrent neural network^15^ or on graph representations placed into a feedforward neural network^16^. The use of graph convolutions allows for more graph nodes to account for adjacent nodes, theoretically improving overall generalisability.

Outside of classification advances and magnitude-level speed improvements, the model is able to explain through its decoded latent space why one molecule is likely to exhibit biological and chemical properties over another, an initiative that is exceedingly challenging for blackbox medical models^26^. The model’s latent space can be exploited to learn in an empirical manner why one type of molecule exhibits a certain biological property more than another as demonstrated by the carcinogens optimisation. The model can also be used to find the intermediate molecules between the two, which can be selected as reference molecules in high-throughput assays, and may enable a better empirical understanding of molecular motifs and their respective biological mechanisms of action. Latent space explainability can also be used to spearhead areas of new research, for example, in using the model to find chemical motifs that can lead to antagonism or agonism against a specific target. Due to the fundamental continuous nature of the latent space, the model can also be used to predict assay results in complex experiments involving more than one mechanism of action. In the generative field, current molecules generated by other generative models are often quite unrealistic, with molecules often being reconstructed from SMILES or some other format^27^. The method proposed, however, overcomes this in a natural way by directly comparing latent spaces or FPs and returning the closest molecule from the search library. This has the added advantage of being able to utilise tailored libraries in which the generated molecules are directly purchasable or synthesisable. Moreover, the use of the latent space potentiation method allows for significant narrowing of molecules, in which a standard drug virtual screen can be performed only on neighbouring molecules, most of which have structural similarities and ideally chemical similarities to the properties which are favoured.

The authors acknowledge three limiting factors of this study: (1) the fingerprints/latent space reverse translation into the original molecules is dependent on the size of the database. Abstract molecules which are not in said database cannot be decoded with high fidelity, with neighbouring approximations being generated; (2) the ZINC clustered database does not contain any transition metal. In model evaluation, unknown atoms are converted to “dummy atoms” to ensure continuity of the graph; (3) the prediction of properties vital to ligand-based drug discovery from potentiating the latent space is not guaranteed to be of high accuracy in every case due to model limitations.

The limiting factors can, however, be mitigated using the corresponding steps: (1) future work can be performed in using generative deep learning models to reconstruct molecules from the fingerprints and/or latent space or larger molecular reference databases can be deployed. (2) Random permutation of molecules in the ZINC database to add transition compounds in a chemically feasible way can be considered, to better train the variational encoder to recognise them; (3) virtual screening of molecules, e.g. through direct molecular docking, that are near the latent space would be necessary instead of using the potentiated molecule directly, owing to lack of fidelity in molecule reconstruction. This is in contrast to the current brute-force technique of screening all the molecules in the dataset, with this method narrowing down the search space.

With all limitations considered, two pipelines can hence be proposed in future CADE: one, to perform a brute force screen of a library using the variational graph encoder, subsequently selecting the top 10% of desired molecules and do further virtual screening. Two, to potentiate the latent space with desired parameters, and extract all neighbouring molecules within a certain latent Euclidean distance or fingerprint distance, and then perform regular virtual screening. The latter of which risks missing potential molecules, however requires an even much lower amount of computational expense. Further work remains to be done in exploring a larger plethora of machine learning models including hyperparameter optimisation and autoML selection, which have been used in other fields such as neuroradiology^28^, which are in the works together with the mitigating factors discussed.

In summary, this work has shown that the latent space of variational graph encoders as described is of surprisingly versatile nature and can be used to predict the properties of highly diverse datasets. Further work will involve the mitigating strategies of limiting factors and the application of the algorithm in a drug discovery pipeline including consequent experimental validation.

## 5. Methods

### 5.1 Clustering of ZINC molecules

**~**690,000,000 ZINC^29^ molecules were split on their tranches with approximately 1,000,000 molecules per segment, and each segment clustered into 10,000 clusters using Morgan2 FPs with 256 bits using the faiss_kmeans library by Pat Walters (https://github.com/PatWalters/faiss_kmeans). The cluster centroids are used for downstream training of the variational graph encoder.

### 5.2 Generation of molecular descriptors and fingerprints

MACCS and Morgan fingerprints were generated using RDKit 2022.03.5, with Morgan fingerprints having a radius of 4 and 512 bits. Mordred descriptors generated using the Mordred 1.2.0 library^30^. For Mordred descriptors in which the output is a non-float number, or is otherwise undefined, it is set to a value of zero.

### 5.3 Graph node and edge generation

SMILES strings are converted into a graph representation where the atoms and the bonds between the atoms are represented by nodes and edges, respectively. In chemical species that have ionic character, the largest ion is chosen. Nodes are represented by a matrix of *N* vectors where *N* is the amount of atoms in the molecule or ion and each vector is a one-hot encoding of the type of the atom. Edges are represented as an adjacency matrix between connected nodes, and a one-hot vector for each edge encoding the bond type. Types of atoms that can be encoded are: Al, As, B, Be, Br, C, Ca, Cl, Co, Cu, F, Fe, H, I, Ir, K, Li, Mg, N, Na, O, Os, P, Pt, Re, Rh, Ru, S, Sb, Se, Si, Te, V and Zn. Atoms which belong to none of the available atom types are encoded as “dummy” atoms. Similarly, types of bonds that can be encoded are: single, double, triple, and aromatic bonds.

### 5.4 Training of the variational graph encoder and architecture

The model was written in Python 3.10 using the Spektral 1.2.0 library using the TensorFlow 2.10.0 backend. The full model is disclosed with its trained weights in the available GitHub repository. In essence, for the first part of the encoder, a masking layer is first applied, followed by a precondition layer with 16 neurons per node. Three layers of edge-conditioned convolutions (ECConv) are then applied, with 32 features per node per layer. A global sum pool is then applied before passing through a feedforward layer with 256 neurons. Subsequently, the tensor is flattened and the values fed into another feedforward layer with 256 neurons. Lastly, a sampling layer is added with 64 latents as per standard variational autoencoders, giving the latent space. The KL loss is also calculated at this step and multiplied by a parameter, β_KL_, which is defined as such:

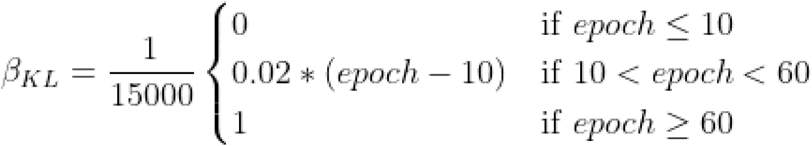

Subsequently, the latent space is fed through the second part of the variational graph encoder, similar to an autoencoder’s decoder segment, consisting of a feedforward layer with 256 neurons. The output of the second half of the encoder is 679 binary FPs achieved with a sigmoid activation function, and 1,613 Mordred regression descriptors with no activation function. Activation functions for the rest of the model unless otherwise specified are leaky rectified linear unit (ReLU) with α = 0.05. The loss of the binary FPs is defined as binary cross-entropy, whereas loss for the regression part was defined as Huber loss with δ = 0.5. The KL loss, binary cross-entropy loss and Huber losses are then summed in 1: 1: 1 ratio and minimised accordingly using gradient descent with a batch size of 64. The model was trained using the Adam optimiser, with a starting learning rate = 2e-4, β_1_ = 0.9, β_2_ = 0.999, and ε = 1e-8. Learning rate was decayed throughout the training using an inverse time schedule. Training was performed on 100 epochs, with 650,000 clustered molecules from the ZINC dataset in each epoch. The remaining 49,999 molecules were sequestered and used as a test set to generate model metrics after training. The model is illustrated in Supplementary Fig. 1.

### 5.5 Surrogate model training and evaluation

Surrogate models were created using either TensorFlow or the Scikit-Learn 1.1.2 library. For extra tree classifiers, 512 estimators were used, with log_2_(x) number of features and no limit to tree depth, where x is the number of features. For extra tree regressors, 2048 estimators were used, similarly with log_2_(x) number of features and no limit to tree depth. Support vector regressors rely on the NuSVR^31^ algorithm with ν = 0.5 and error term C = 0.5. Standard normalisation was applied with outlier values more than the 90th percentile and less than the 10th percentile removed before training for the extra trees models, and outliers were not removed for the support vector regression method in training. Models in which multiple molecules were required had latent space concatenated before input. The TWOSIDES polypharmacy set’s raw labels were classified using a feedforward neural network of 512 × 5 layers, and an output of the number of raw labels. Evaluations on datasets were performed using fivefold cross-validations for all cases unless otherwise specified, with the dataset initially randomly shuffled to prevent bias in any cross-validation segments. Other surrogate models evaluated include XGBoost from the XGBoost 1.6.2 library, random forests, gradient boosting, support vector machines, K-neighbours and Gaussian processes. Fivefold cross-validation using a random split was used unless otherwise specified for all surrogate models.

### 5.6 Databases and datasets used for surrogate model training

Datasets were downloaded from the TDCommons, ChEMBL and Tox21 databases. Datasets used from the TDCommons database include the hERG^32^, CYP P450^33–35^, carcinogens^22,35^, blood brain barrier^36^, mutagenicity^37^, human intestinal absorption^38^, drug-induced liver injury^39^, skin reaction^40,41^, LD_50_^42^, steady state volume distribution^43^, plasma protein binding rate^44^, half life^45^, hepatocyte and microsomal clearance^44,46^, bioavailability^47^, lipophilicity^44,48^, solubility^49^, solvation free energy^48,50^, skin reaction^40^, anti-HIV virus activity^48^, anti-SARS-CoV-2^51^, activity against SARS-CoV-2 3CL protease^52^, TWOSIDES polypharmacy^53^ and DDI datasets^54,55^. Entries in a single Tox21^18^ dataset that had duplicate SMILES codes were removed to prevent data leakage across cross-validation sets. ChEMBL sets used include: ChEMBL1614672, ChEMBL2093836, and ChEMBL1614236.

### 5.7 Reclassification of TWOSIDES polypharmacy dataset

The TWOSIDES polypharmacy dataset was reclassified into 26 categories according to the International Classification of Diseases 11th Revision (ICD-11)^56^. Ambiguous labels or diseases not found in the database were classified under “others”. The reclassification of the side effects can be found in Supplementary File 2.

### 5.8 Ligand-based drug discovery surrogate models training

X-ray crystallography-resolved proteins with PDB IDs: 2QFT, 2XFH, 4EJW, 5TUL, and 7B3C were downloaded from the RCSB PDB database and Gasteiger charges added using ADFR Suite^57^ after the removal of non-protein and non co-ligand molecules (i.e. water, ions used in protein crystallisation). Docking pockets were identified using existing native ligands and 50,000 randomly selected molecules from the ChemBridge Express and Core libraries (http://chembridge.com) were docked with either QuickVina2.1^58^ or Gnina^59^. In both docking cases a uniform box size of 20 Å was used. Rescoring functions, namely OnionNet-SFCT^60^, RTMScore^61^ and DeepRMSD^62^ were applied on the QuickVina2.1 docked poses. Fivefold cross-validation as described in section 6.5 was then used to evaluate model efficacy.

### 5.9 Hardware for speed evaluation

An Nvidia RTX^™^ 3090 was used for GPU measurements of speed. The CPU used is an Advanced Micro Devices Ryzen^™^ 5950X. DDR4 RAM was employed for all model calculations pertinent to speed benchmarking. Speed tests were performed uniformly on this set of hardware running Ubuntu 22.04.1 LTS (GNU/Linux 5.15.0-56-generic x86_64) with no other foreground programs running when benchmarking was conducted.

### 5.10 Simulated annealing for drug potentiation

Drug potentiation simulated annealing was performed with an annealing α = 0.99 for 10,000 iterations and a starting temperature β = 1, as described by Pincus *et al.*, 1970^63^. Each step was performed by adding Gaussian noise with a standard deviation of 0.01 and mean of 0 to the latent space of the last accepted iteration of the Markov chain. The initial latent space is a Gaussian vector with standard deviation of 0.5 and mean of 0. Reward functions are defined with the following equation:

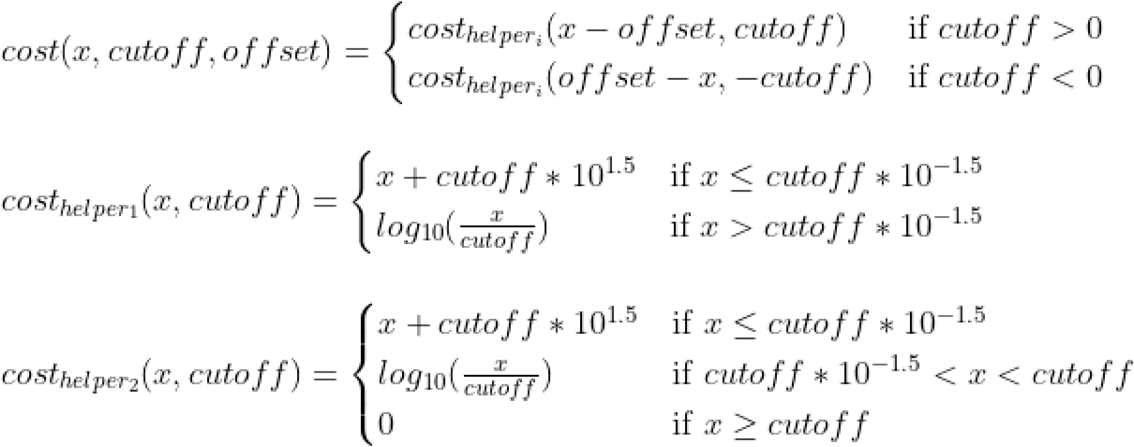

The sum of all the cost functions is then used as the reward function for the simulated annealing, with a higher value being more desirable.

### 5.11 Molecule decoding from latent space or output fingerprints

Latent space was decoded through Euclidean distance with a library of all clustered ZINC molecules unless otherwise specified. All clustered molecules in the library were individually evaluated in the model and the Euclidean distance measured between the reference molecule’s latent space, with the molecule having the smallest Euclidean distance being chosen as the best match. When the fingerprint was decoded, a sum of the absolute error between model-generated fingerprints and reference library was used, with the reference molecule with the best match returned.

## 6. Acknowledgements

The authors thank Tan Lai Heng and Saxena Shikhar for their comments in the initial phase of the work and Thomas Larry Dawson, Jr. for his continued support.

## 7. Code Availability

Code is available at https://github.com/Chokyotager/NotYetAnotherNightshade.

## 9. Funding

This work is supported by the Singapore Ministry of Education (MOE) Tier 1 grant RG27/21 and Tier 2 grant MOE-T2EP30120-0007. Hilbert Lam Yuen In is also supported by funding from the Agency for Science, Technology and Research (A*STAR) and A*STAR BMRC EDB IAF-PP grants (H17/01/a0/004, Skin Research Institute of Singapore, and H18/01a0/016, Asian Skin Microbiome Program). The computational work for this article was partially performed on resources of the National Supercomputing Centre, Singapore (https://www.nscc.sg). SCELSE is funded by Singapore’s National Research Foundation, the Ministry of Education, Nanyang Technological University (NTU), and the National University of Singapore (NUS), and is hosted by NTU in partnership with NUS.

---

**Supplementary Figure 1.**
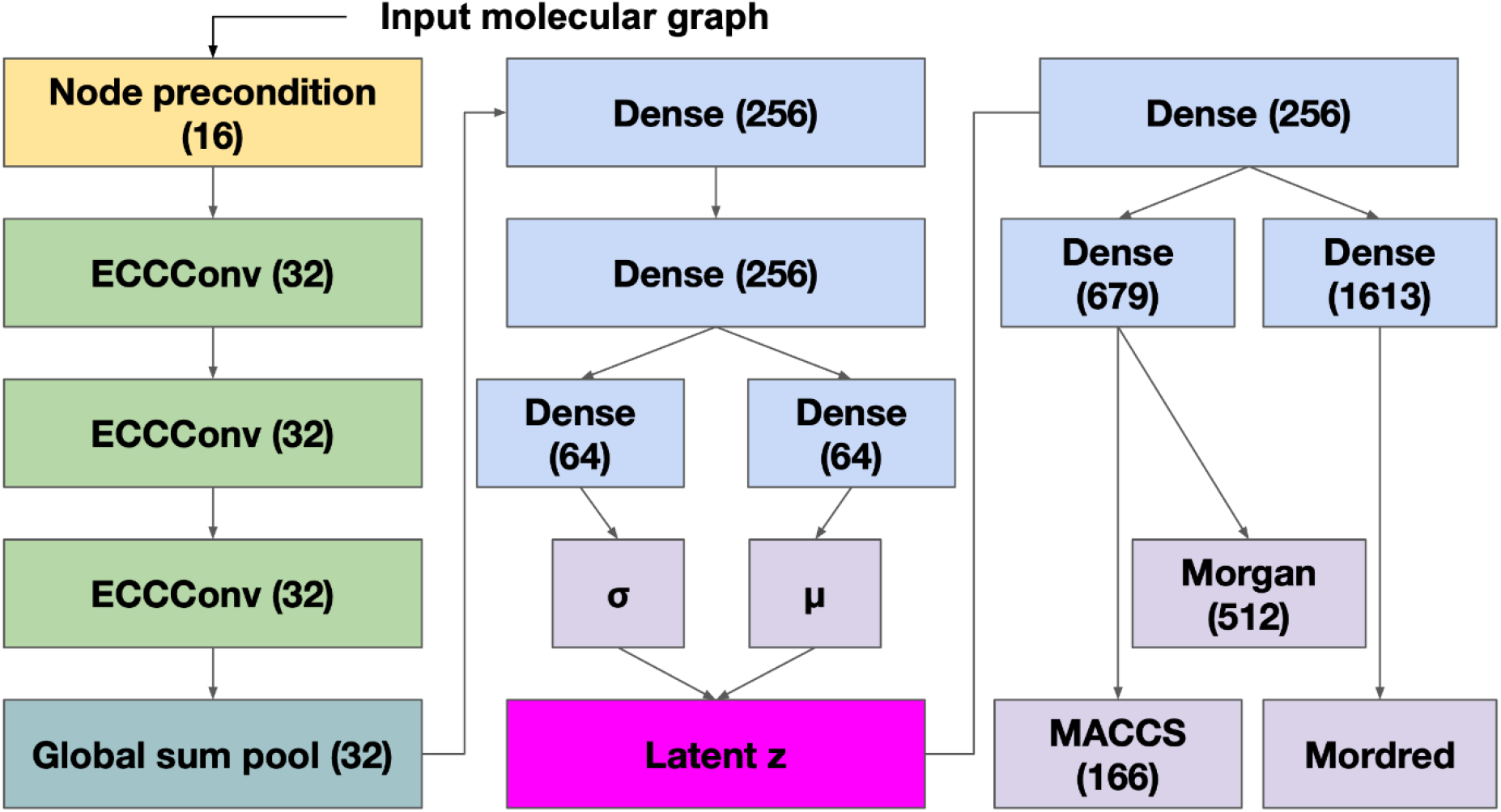
Model architecture overview of variational graph encoder. Three losses are added to train the model - Kullback-Leibler divergence loss, binary cross-entropy for MACCS and Morgan4 FPs, and Huber loss for Mordred descriptors.

**Supplementary Figure 2.**
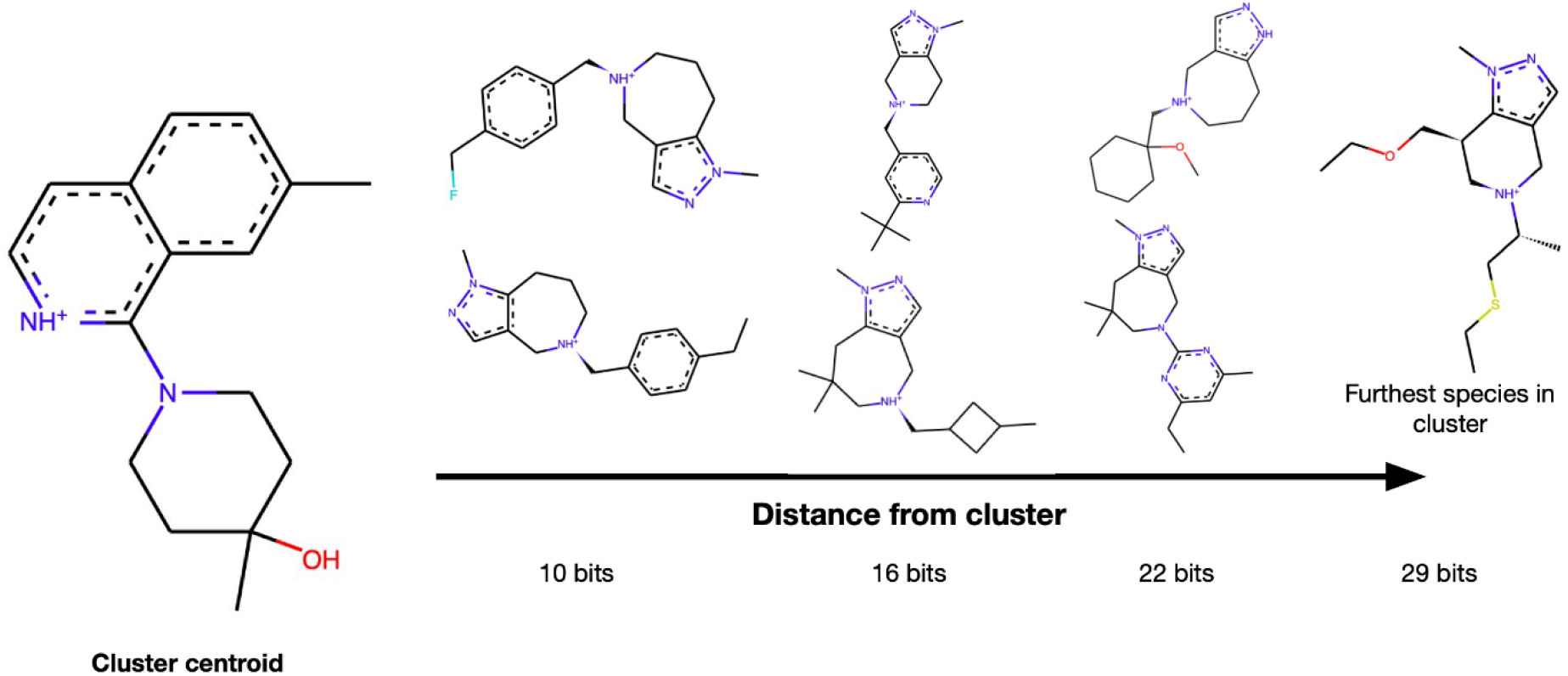
Clusters of molecules in ZINC show overall structural similarity. Molecules were clustered using the faiss_kmeans library, with each 10,000 molecules having approximately one cluster. Cluster centroid shown is one of 700,000, with other molecules shown being part of the same cluster and their FP distance shown.

**Supplementary Figure 3.**
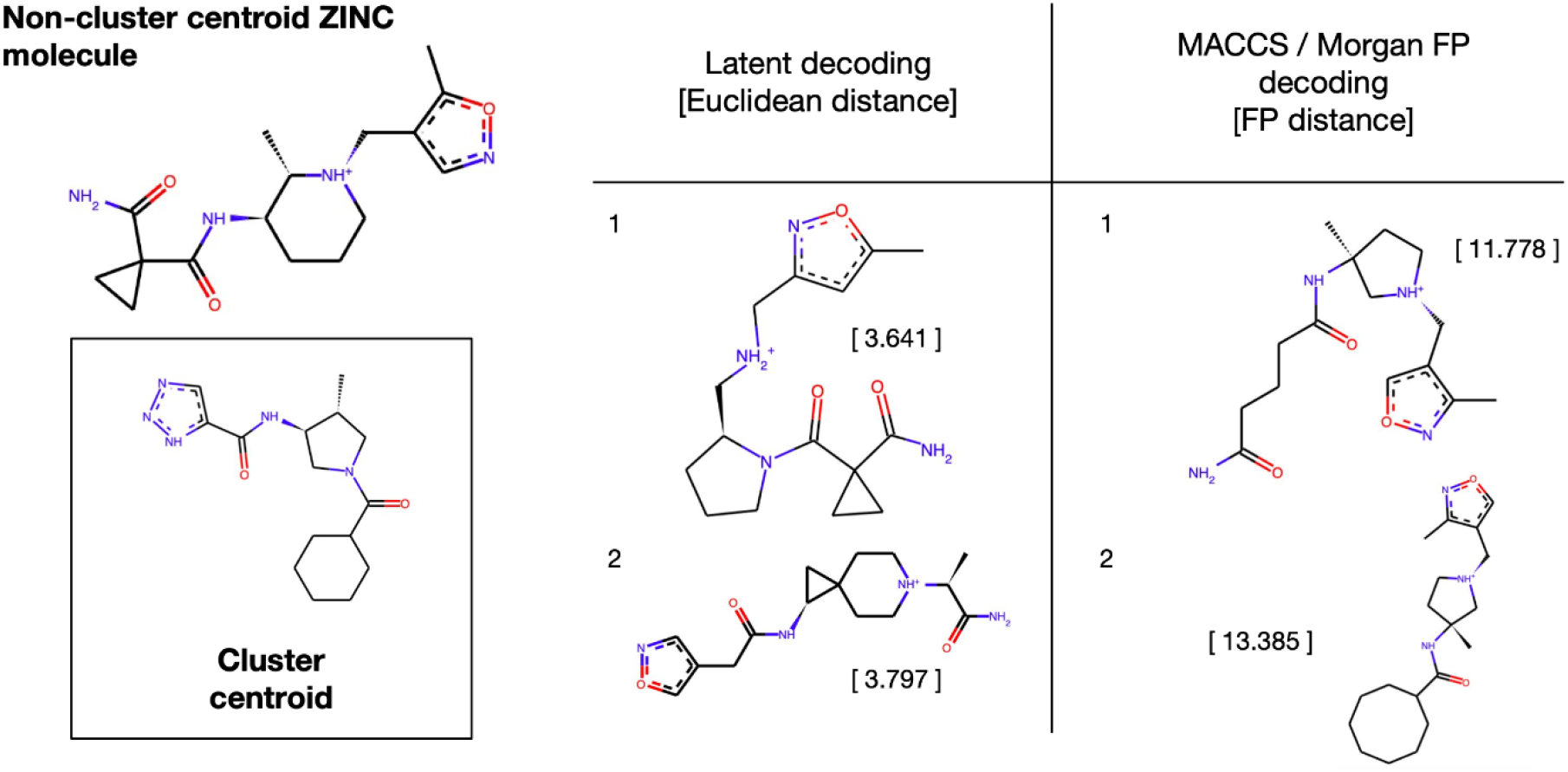
Decoding through latent space matches generally shows better visual fidelity than through FP matching. Non-cluster centroid ZINC molecule was parsed through the variational graph encoder and was then decoded either from the latent space or from FP distance. Scores shown in squared bracket for latent decoding indicate the displayed molecule’s Euclidean distance from the ZINC molecule, and FP distance is shown when FP was used for decoding.

**Supplementary Figure 4.**
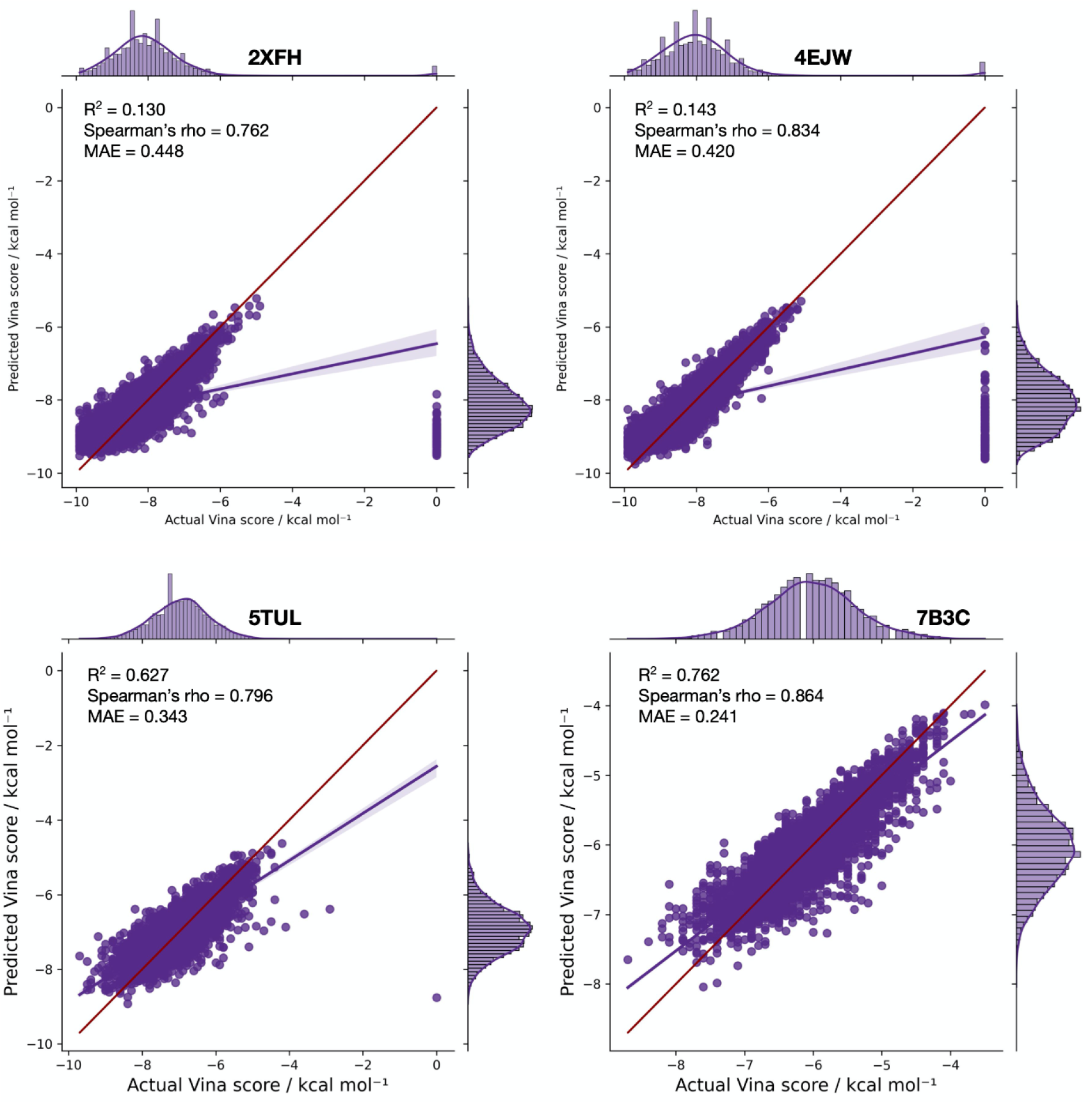
Surrogate regression Vina models show strong predictive power but do not generalise well to multimodal sets. Molecules were docked to respective proteins using QuickVina2.1. Values shown are raw values versus those predicted by the surrogate NuSVM model on ~45,000 ligands, and a 10% (~5,000) segregated testing set versus ground truth scores.

**Supplementary File 1. Scores and comparisons of all surrogate models. [Excel]**

**Supplementary File 2. TWOSIDES polypharmacy labels as reclassified using the ICD-11 as reference. [TSV]**

**Clustered molecules and respective centroids are available upon request (18 GB).**

## Notes

### Competing Interest Statement

The authors have declared no competing interest.

### Summary of Updates

Added some additional funding information. Supplementary figures 2 and 3 have been moved to 1 and 2 respectively, with an additional statement indicating data availability for the original supplementary figure 1.

## References

1 Bohacek, R. S., McMartin, C. & Guida, W. C. The art and practice of structure-based drug design: a molecular modeling perspective. Med Res Rev 16, 3–50, doi:10.1002/(SICI)1098-1128(199601)16:1<3::AID-MED1>3.0.CO;2-6 (1996).

2 Hutchinson, L. & Kirk, R. High drug attrition rates--where are we going wrong? Nat Rev Clin Oncol 8, 189–190, doi:10.1038/nrclinonc.2011.34 (2011).

3 Wouters, O. J., McKee, M. & Luyten, J. Estimated Research and Development Investment Needed to Bring a New Medicine to Market, 2009-2018. JAMA 323, 844–853, doi:10.1001/jama.2020.1166 (2020).

4 Baig, M. H., Ahmad, K., Rabbani, G., Danishuddin, M. & Choi, I. Computer Aided Drug Design and its Application to the Development of Potential Drugs for Neurodegenerative Disorders. Curr Neuropharmacol 16, 740–748, doi:10.2174/1570159X15666171016163510 (2018).

5 Liu, T. et al. Applying high-performance computing in drug discovery and molecular simulation. Natl Sci Rev 3, 49–63, doi:10.1093/nsr/nww003 (2016).

6 Sun, D., Gao, W., Hu, H. & Zhou, S. Why 90% of clinical drug development fails and how to improve it? Acta Pharm Sin B 12, 3049–3062, doi:10.1016/j.apsb.2022.02.002 (2022).

7 Tornio, A., Filppula, A. M., Niemi, M. & Backman, J. T. Clinical Studies on Drug-Drug Interactions Involving Metabolism and Transport: Methodology, Pitfalls, and Interpretation. Clin Pharmacol Ther 105, 1345–1361, doi:10.1002/cpt.1435 (2019).

8 Wang, J. Comprehensive assessment of ADMET risks in drug discovery. Curr Pharm Des 15, 2195–2219, doi:10.2174/138161209788682514 (2009).

9 Kwon, S., Bae, H., Jo, J. & Yoon, S. Comprehensive ensemble in QSAR prediction for drug discovery. BMC Bioinformatics 20, 521, doi:10.1186/s12859-019-3135-4 (2019).

10 Wang, J. & Skolnik, S. Recent advances in physicochemical and ADMET profiling in drug discovery. Chem Biodivers 6, 1887–1899, doi:10.1002/cbdv.200900117 (2009).

11 Wu, F. et al. Computational Approaches in Preclinical Studies on Drug Discovery and Development. Front Chem 8, 726, doi:10.3389/fchem.2020.00726 (2020).

12 Diederik P Kingma, M. W. Auto-Encoding Variational Bayes. arXiv (2013).

13 Li, Y. et al. Generative deep learning enables the discovery of a potent and selective RIPK1 inhibitor. Nat Commun 13, 6891, doi:10.1038/s41467-022-34692-w (2022).

14 Yang, L. et al. Transformer-Based Generative Model Accelerating the Development of Novel BRAF Inhibitors. ACS Omega 6, 33864–33873, doi:10.1021/acsomega.1c05145 (2021).

15 Gomez-Bombarelli, R. et al. Automatic Chemical Design Using a Data-Driven Continuous Representation of Molecules. ACS Cent Sci 4, 268–276, doi:10.1021/acscentsci.7b00572 (2018).

16 Lee, M. & Min, K. MGCVAE: Multi-Objective Inverse Design via Molecular Graph Conditional Variational Autoencoder. J Chem Inf Model 62, 2943–2950, doi:10.1021/acs.jcim.2c00487 (2022).

17 Martin Simonovsky, N. K. Dynamic Edge-Conditioned Filters in Convolutional Neural Networks on Graphs. arXiv (2017).

18 Richard, A. M. et al. The Tox21 10K Compound Library: Collaborative Chemistry Advancing Toxicology. Chem Res Toxicol 34, 189–216, doi:10.1021/acs.chemrestox.0c00264 (2021).

19 Huang, K. et al. Artificial intelligence foundation for therapeutic science. Nat Chem Biol 18, 1033–1036, doi:10.1038/s41589-022-01131-2 (2022).

20 Gaulton, A. et al. ChEMBL: a large-scale bioactivity database for drug discovery. Nucleic Acids Res 40, D1100–1107, doi:10.1093/nar/gkr777 (2012).

21 Maia, E. H. B., Assis, L. C., de Oliveira, T. A., da Silva, A. M. & Taranto, A. G. Structure-Based Virtual Screening: From Classical to Artificial Intelligence. Front Chem 8, 343, doi:10.3389/fchem.2020.00343 (2020).

22 Lagunin, A. A., Dearden, J. C., Filimonov, D. A. & Poroikov, V. V. Computer-aided rodent carcinogenicity prediction. Mutat Res 586, 138–146, doi:10.1016/j.mrgentox.2005.06.005 (2005).

23 Hansen, P. & Bichel, J. Carcinogenic effect of sulfonamides. Acta radiol 37, 258–265, doi:10.3109/00016925209139877 (1952).

24 Littlefield, N. A., Sheldon, W. G., Allen, R. & Gaylor, D. W. Chronic toxicity/carcinogenicity studies of sulphamethazine in Fischer 344/N rats: two-generation exposure. Food Chem Toxicol 28, 157–167, doi:10.1016/0278-6915(90)90004-7 (1990).

25 Masumshah, R., Aghdam, R. & Eslahchi, C. A neural network-based method for polypharmacy side effects prediction. BMC Bioinformatics 22, 385, doi:10.1186/s12859-021-04298-y (2021).

26 Price, W. N. Big data and black-box medical algorithms. Sci Transl Med 10, doi:10.1126/scitranslmed.aao5333 (2018).

27 Zeng, X. et al. Deep generative molecular design reshapes drug discovery. Cell Rep Med 3, 100794, doi:10.1016/j.xcrm.2022.100794 (2022).

28 Musigmann, M. et al. Testing the applicability and performance of Auto ML for potential applications in diagnostic neuroradiology. Sci Rep 12, 13648, doi:10.1038/s41598-022-18028-8 (2022).

29 Irwin, J. J. & Shoichet, B. K. ZINC--a free database of commercially available compounds for virtual screening. J Chem Inf Model 45, 177–182, doi:10.1021/ci049714+ (2005).

30 Moriwaki, H., Tian, Y. S., Kawashita, N. & Takagi, T. Mordred: a molecular descriptor calculator. J Cheminform 10, 4, doi:10.1186/s13321-018-0258-y (2018).

31 Platt, J. Probabilistic outputs for support vector machines and comparisons to regularized likelihood methods. (1999).

32 Wang, S. et al. ADMET Evaluation in Drug Discovery. 16. Predicting hERG Blockers by Combining Multiple Pharmacophores and Machine Learning Approaches. Mol Pharm 13, 2855–2866, doi:10.1021/acs.molpharmaceut.6b00471 (2016).

33 Veith, H. et al. Comprehensive characterization of cytochrome P450 isozyme selectivity across chemical libraries. Nat Biotechnol 27, 1050–1055, doi:10.1038/nbt.1581 (2009).

34 Carbon-Mangels, M. & Hutter, M. C. Selecting Relevant Descriptors for Classification by Bayesian Estimates: A Comparison with Decision Trees and Support Vector Machines Approaches for Disparate Data Sets. Mol Inform 30, 885–895, doi:10.1002/minf.201100069 (2011).

35 Cheng, F. et al. admetSAR: a comprehensive source and free tool for assessment of chemical ADMET properties. J Chem Inf Model 52, 3099–3105, doi:10.1021/ci300367a (2012).

36 Martins, I. F., Teixeira, A. L., Pinheiro, L. & Falcao, A. O. A Bayesian approach to in silico blood-brain barrier penetration modeling. J Chem Inf Model 52, 1686–1697, doi:10.1021/ci300124c (2012).

37 Xu, C. et al. In silico prediction of chemical Ames mutagenicity. J Chem Inf Model 52, 2840–2847, doi:10.1021/ci300400a (2012).

38 Hou, T., Wang, J., Zhang, W. & Xu, X. ADME evaluation in drug discovery. 7. Prediction of oral absorption by correlation and classification. J Chem Inf Model 47, 208–218, doi:10.1021/ci600343x (2007).

39 Xu, Y. et al. Deep Learning for Drug-Induced Liver Injury. J Chem Inf Model 55, 2085–2093, doi:10.1021/acs.jcim.5b00238 (2015).

40 Alves, V. M. et al. Predicting chemically-induced skin reactions. Part I: QSAR models of skin sensitization and their application to identify potentially hazardous compounds. Toxicol Appl Pharmacol 284, 262–272, doi:10.1016/j.taap.2014.12.014 (2015).

41 National Institute of Environmental Health Sciences (NIEHS); the Murine Local Lymph Node Assay: a Test Method for Assessing the Allergic Contact Dermatitis Potential of Chemicals/Compounds, report now available. Public Health Service. Fed Regist 64, 14006–14007 (1999).

42 Zhu, H. et al. Quantitative structure-activity relationship modeling of rat acute toxicity by oral exposure. Chem Res Toxicol 22, 1913–1921, doi:10.1021/tx900189p (2009).

43 Lombardo, F. & Jing, Y. In Silico Prediction of Volume of Distribution in Humans. Extensive Data Set and the Exploration of Linear and Nonlinear Methods Coupled with Molecular Interaction Fields Descriptors. J Chem Inf Model 56, 2042–2052, doi:10.1021/acs.jcim.6b00044 (2016).

44 Mark Wenlock, N. T. Experimental in vitro DMPK and physicochemical data on a set of publicly disclosed compounds. doi:10.6019/CHEMBL3301361.

45 Obach, R. S., Lombardo, F. & Waters, N. J. Trend analysis of a database of intravenous pharmacokinetic parameters in humans for 670 drug compounds. Drug Metab Dispos 36, 1385–1405, doi:10.1124/dmd.108.020479 (2008).

46 Di, L. et al. Mechanistic insights from comparing intrinsic clearance values between human liver microsomes and hepatocytes to guide drug design. Eur J Med Chem 57, 441–448, doi:10.1016/j.ejmech.2012.06.043 (2012).

47 Ma, C. Y. et al. Prediction models of human plasma protein binding rate and oral bioavailability derived by using GA-CG-SVM method. J Pharm Biomed Anal 47, 677–682, doi:10.1016/j.jpba.2008.03.023 (2008).

48 Wu, Z. et al. MoleculeNet: a benchmark for molecular machine learning. Chem Sci 9, 513–530, doi:10.1039/c7sc02664a (2018).

49 Sorkun, M. C., Khetan, A. & Er, S. AqSolDB, a curated reference set of aqueous solubility and 2D descriptors for a diverse set of compounds. Sci Data 6, 143, doi:10.1038/s41597-019-0151-1 (2019).

50 Mobley, D. L. & Guthrie, J. P. FreeSolv: a database of experimental and calculated hydration free energies, with input files. J Comput Aided Mol Des 28, 711–720, doi:10.1007/s10822-014-9747-x (2014).

51 Touret, F. et al. In vitro screening of a FDA approved chemical library reveals potential inhibitors of SARS-CoV-2 replication. Sci Rep 10, 13093, doi:10.1038/s41598-020-70143-6 (2020).

52 diamond. Main protease structure and XChem fragment screen. (2020).

53 Tatonetti, N. P., Ye, P. P., Daneshjou, R. & Altman, R. B. Data-driven prediction of drug effects and interactions. Sci Transl Med 4, 125ra131, doi:10.1126/scitranslmed.3003377 (2012).

54 Ryu, J. Y., Kim, H. U. & Lee, S. Y. Deep learning improves prediction of drug-drug and drug-food interactions. Proc Natl Acad Sci U S A 115, E4304–E4311, doi:10.1073/pnas.1803294115 (2018).

55 Wishart, D. S. et al. DrugBank 5.0: a major update to the DrugBank database for 2018. Nucleic Acids Res 46, D1074–D1082, doi:10.1093/nar/gkx1037 (2018).

56 Organization, W. H. International Classification of Diseases, Eleventh Revision (ICD-11). (2019).

57 Ravindranath, P. A., Forli, S., Goodsell, D. S., Olson, A. J. & Sanner, M. F. AutoDockFR: Advances in Protein-Ligand Docking with Explicitly Specified Binding Site Flexibility. PLoS Comput Biol 11, e1004586, doi:10.1371/journal.pcbi.1004586 (2015).

58 Alhossary, A., Handoko, S. D., Mu, Y. & Kwoh, C. K. Fast, accurate, and reliable molecular docking with QuickVina 2. Bioinformatics 31, 2214–2216, doi:10.1093/bioinformatics/btv082 (2015).

59 McNutt, A. T. et al. GNINA 1.0: molecular docking with deep learning. J Cheminform 13, 43, doi:10.1186/s13321-021-00522-2 (2021).

60 Zheng, L. et al. Improving protein-ligand docking and screening accuracies by incorporating a scoring function correction term. Brief Bioinform 23, doi:10.1093/bib/bbac051 (2022).

61 Shen, C. et al. Boosting Protein-Ligand Binding Pose Prediction and Virtual Screening Based on Residue-Atom Distance Likelihood Potential and Graph Transformer. J Med Chem 65, 10691–10706, doi:10.1021/acs.jmedchem.2c00991 (2022).

62 Wang, Z. et al. A fully differentiable ligand pose optimization framework guided by deep learning and a traditional scoring function. Brief Bioinform, doi:10.1093/bib/bbac520 (2022).

63 Pincus, M. Letter to the Editor—A Monte Carlo Method for the Approximate Solution of Certain Types of Constrained Optimization Problems. Operations Research, doi:https://doi.org/10.1287/opre.18.6.1225 (1970).

